# Zebrafish *armc9* mutant reveals a link between ARM-Type fold domain loss, inflammation, and spinal curvature

**DOI:** 10.1101/2025.04.02.646874

**Authors:** Anastassia Rubilar-Fajardo, Carlos Muñoz-Montecinos, Gustavo Esperguel-Toledo, Sofía Carrasco, Anguela Castro, Paloma Alvarado, Amy Kugath, Raúl Araya-Secchi, Miguel Allende, Sylvain Marcellini, Marcelo González-Ortiz, Juan Román, Sebastián Boltaña, Felipe Aguilera, Timothy Cooper, Jean Cooper, Keith Cheng, Teresa Caprile, Mary C. Mullins, Ricardo Fuentes

**Author notes:** These authors contributed equally (Anastassia Rubilar-Fajardo, Carlos Muñoz-Montecinos).

## Abstract

The vertebral column (VC) defines the central axis of vertebrates. Among its malformations, adolescent idiopathic scoliosis (AIS) is the most common, characterized by lateral spine curvature without congenital abnormalities. While its etiology remains elusive, genetic studies in zebrafish have linked AIS to skeletogenesis, cilia structure, cerebrospinal fluid flow, and inflammation. In this study, we characterized the zebrafish *armc9* mutant, exhibiting AIS-like curvature without vertebral malformations, subcommissural organ, or Reissner fiber structure disruptions. We also detected an inflammatory response at the curvature origin. Structural bioinformatics and molecular docking analyses suggest that the mutant Armc9 protein partially lacks the functional ARM-type fold domain, which is crucial for its interaction with Togaram1, revealing that both proteins are relevant for ciliary signaling and spine curvature. Phylogenetic analyses indicate that the ARMC gene family encompasses 12 members, highlighting that the Armc9 gene has deep evolutionary roots. Our results demonstrate that the zebrafish *armc9* mutant accurately represents the AIS phenotype, suggesting that the loss of Armc9’s C-terminal ARM-type fold domain could affect the ciliary integrity and contribute to spine curvature by altering the skeletogenic environment. Taken together, this study proposes that the zebrafish *armc9* mutant offers a unique opportunity to understand the etiology of AIS and provides new insights into the molecular interaction networks of Armc9 in ciliary signaling.

## Introduction

The vertebral column (VC), or spine, is a defining feature of vertebrate animals, forming the central axis of the skeleton and providing structural support, spinal cord protection, and anchorage for other bones (1). Abnormal VC development can lead to congenital deformities, including kyphosis (excessive thoracic curvature), lordosis (abnormal lumbar curvature), and scoliosis (a complex three-dimensional deformity of the spine and trunk) (2). Scoliosis is the most common VC deformity in children and adolescents, with adolescent idiopathic scoliosis (AIS) accounting for 80% of cases worldwide (3). AIS is defined by a lateral spinal curvature exceeding 10 degrees in a standing patient, often accompanied by rotational defects (4). While AIS is generally not a progressive condition, scoliosis can cause various problems depending on the degree of the curvature. Despite extensive research on AIS morphogenesis and ongoing efforts to develop therapeutic strategies, the cellular and molecular mechanisms underlying VC curvature development during vertebrate post-embryonic stages remain poorly understood.

Advances in genetics and high-throughput sequencing in animal models have facilitated the identification of genes implicated in AIS etiology (5). Expanding this knowledge and uncovering the genetic basis of VC development is essential for improving AIS diagnosis, prognosis, and treatment. However, while animal models have successfully replicated spinal curvature resembling AIS, they have not fully recapitulated its etiopathogenesis, limiting insights into its underlying causes (6). This underscores the need for alternative experimental strategies and model systems that can faithfully mirror AIS-specific characteristics.

Interestingly, unlike mammals, certain vertebrate species, such as fish, naturally develop scoliosis. Scoliosis-like spinal curvatures have been observed in several teleost species, including guppies, medaka, and zebrafish (*Danio rerio*) (5, 7, 8). As a vertebrate model, zebrafish exhibit structural similarities to the human VC, including vertebrae with intervertebral discs, rib associations, and a central notochord (9). Indeed, the zebrafish model has advanced AIS research, revealing the relevant role of cilia and cerebrospinal fluid (CSF) (5, 10). This makes zebrafish an invaluable system for studying AIS pathogenesis, particularly in the context of ciliary function and CSF dynamics.

Cilia are elongated structures that protrude from the cell membrane into the extracellular space and play crucial roles in cellular signaling and fluid movement. They are broadly classified into motile cilia, which drive fluid flow and create signaling gradients, and primary cilia, which mediate signal transduction to regulate intracellular processes (11, 12). Defects in ciliary structure or function underlie ciliopathies, a group of disorders affecting multiple organs, including the brain, kidneys, and skeleton (13). Recent studies have highlighted the importance of ciliary homeostasis in VC development, with disruptions in both primary and motile cilia linked to scoliosis (9, 14, 15). In this context, zebrafish has significantly advanced our understanding of AIS, particularly by revealing the role of cilia in maintaining body axis straightness. For instance, mutations in the *ptk7* gene, which disrupt ependymal cell cilia, result in scoliosis-like curvatures (16). Similarly, other ciliary gene mutations produce axial curvature reminiscent of human ciliopathies (17). Another relevant component with an intricate relationship with cilia is the Reissner fiber (RF). This is a dynamic structure formed by the subcommissural organ (SCO)-secreted and highly conserved glycoprotein SCO-spondin (18). RF plays a crucial role in early body axis linearity, and its defective formation has been linked to scoliosis development in zebrafish (19–21). Moreover, scoliosis in zebrafish has been associated with increased inflammatory signaling, particularly involving TNF-α, further implicating immune responses in AIS pathogenesis (5, 22, 23).

Primary cilia also regulate key developmental pathways, including Hedgehog, Wnt, and TGF-β/BMP signaling, which control skeletal development (24). One essential ciliary protein, Centrosomal protein 290 (Cep290), is involved in ciliogenesis and has been implicated in multiple human ciliopathies, such as Joubert syndrome (25). Zebrafish *cep290* mutants recapitulate human ciliopathies, displaying axial curvature, cerebellar defects, kidney cysts, and vision impairments (26–28). Overall, ciliary function, both primary and motile, is essential for the development and proper formation of the axial axis. Dysregulation in signaling pathways such as Hedgehog or CSF flow are the first processes related to the development of pathological phenotypes such as AIS, thus paving the way for understanding the etiology of this condition.

Among the genes associated with AIS is *armc9*, which encodes the Armadillo repeat-containing protein 9 (Armc9). Whole-exome sequencing linked *armc9* mutations to Joubert syndrome (29). Like other proteins previously associated with this syndrome, Armc9 is localized in cilia. Specifically, in the centriole and basal body of primary cilia, and it has been shown to migrate to the distal region in response to Hedgehog signaling (29, 30), where it interacts with known ciliopathy-associated proteins such as CEP290, CEP104, CSPP1, CCDC66, and TOGARAM1 (31, 32). Armc9-deficient cells exhibit shortened cilia, reduced Hedgehog pathway regulators (GLI2 and GLI3), and altered tubulin modifications, suggesting a role in ciliary protein trafficking (30). Additionally, splicing variants of *armc9* have been linked to developmental disorders, including mental retardation, ptosis, and polydactyly (33). Zebrafish *armc9* mutants have provided new insights into ciliary function and AIS. These mutants exhibit hallmark ciliopathy features, retinal dystrophy, reduced ventricular cilia, and axial curvature, resembling AIS (29). However, the underlying morphogenetic processes contributing to this phenotype remain unclear. Moreover, previous studies on zebrafish *armc9* mutants have focused primarily on adult individuals, leaving early developmental aspects of AIS etiology unexplored.

Here, we present a phenotypic, molecular, and evolutionary characterization of the zebrafish *armc9* mutant displaying severe scoliosis. Our findings indicate that the observed spinal curvature is not caused by defects in bone structure and/or RF formation, but associated to an inflammatory response. At the molecular level, zebrafish Armc9 protein would interact with Tog1 through its ARM-type fold domain, revealing key functional residues responsible for Armc9-Tog1 interaction. Additionally, a phylogenetic analysis of 148 ARMC family proteins across multiple animal species identified 12 phylogenetic groups corresponding to known family members and that Armc9 has its evolutionary origin at the dawn of the Metazoan. Collectively, our results suggest the involvement of a previously unrecognized mechanisms in scoliosis pathogenesis, expanding current models of AIS etiology. Furthermore, our findings contribute to the functional characterization of Armc9-Tog1 interactions and the evolutionary origins of ARMC family proteins in vertebrates.

## Results

### The armc9^sa29834^ mutant phenotype resembles an AIS-like spinal curvature

The *armc9^sa29834^* mutant was generated and isolated through a genetic screen using TILLING technology (34). *armc9* mutant fish exhibit a significant reduction in body length along the anterior-to-posterior axis, accompanied by a pronounced dorsal hump that is absent in the wild-type fish (Fig. 1A, B). To assess the extent of the axial curvature and skeletal abnormalities, we performed X-ray imaging for a detailed examination of the VC (Fig. 1A). Skeletal analysis revealed that the wild-type VC maintains a straight and orderly vertebrate-based alignment, consistent with normal axial development (Fig. 1A). In contrast, *armc9* mutants display severe curvature of the VC, particularly along the anterior-to-posterior axis, indicating a substantial skeletal malformation. These imaging techniques highlights the severity of the axial deformities in the mutant phenotype, suggesting that *armc9* plays a crucial role in maintaining VC structure, axial alignment, and function.

**Figure 1.**
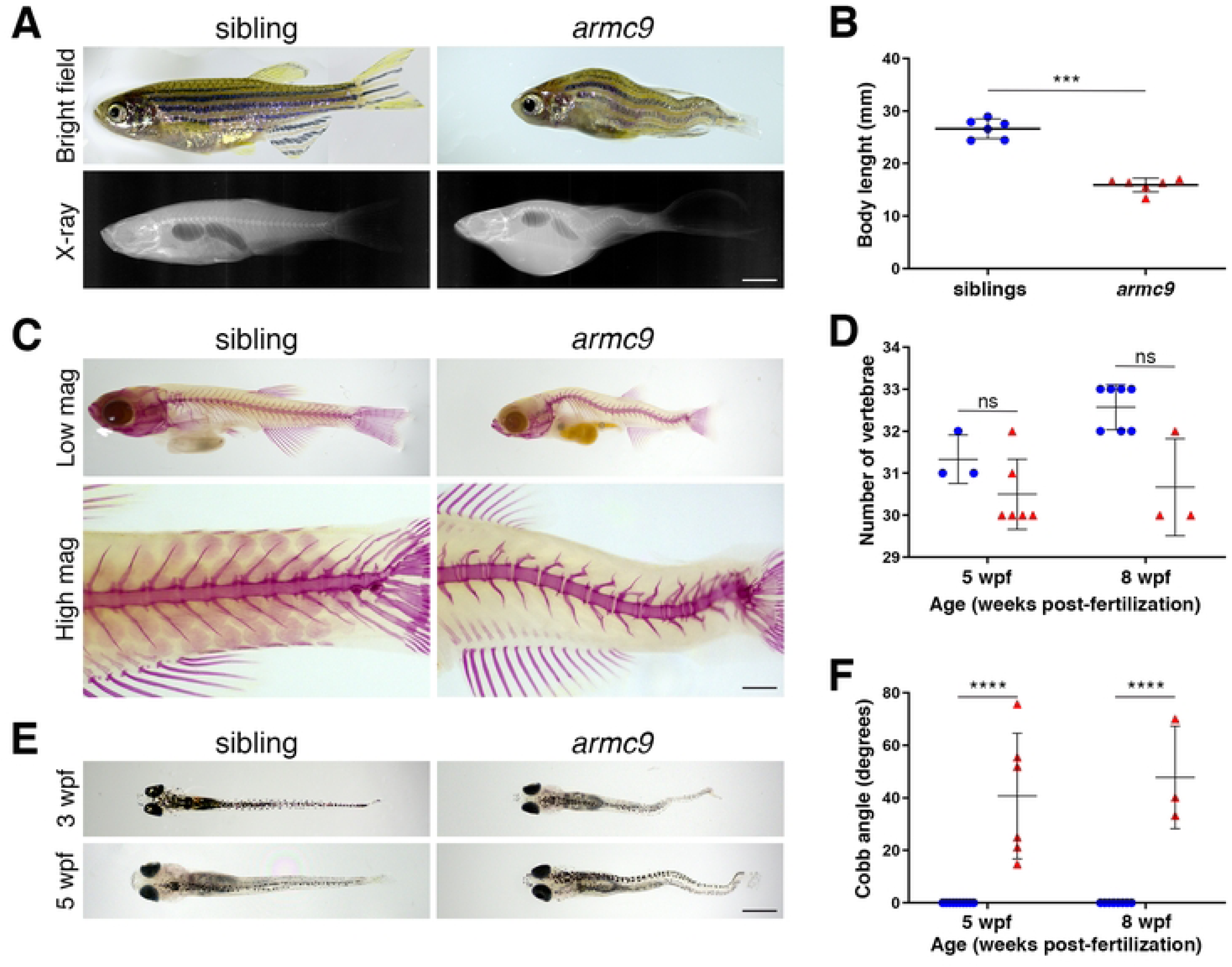
The *armc9^sa29834^* mutant phenotype. **A.** Top row: Lateral view of a representative adult wild-type sibling and *armc9^sa29834^* fish. Mutants exhibit a remarkable reduction in the anterior-to-posterior axis compared to wild-type adults, along with a pronounced dorsal curvature. Bottom row: X-ray images reveal that body axis curvature is caused by a scoliosis-like deformation of the vertebral column (VC). **B.** Scatter plot of body length measurements along the anterior-to-posterior axis in adult wild-type sibling and *armc9^sa29834^* fish. The average body length in mutants was significantly different from that of wild-type adults (WT= 26.63 mm, *armc9*= 15.91 mm). Sample sizes: wild type (n= 6 adults) and *armc9* (n= 6 adults). Data are presented as mean ± SEM. \*\*\**p*<0.05 (Student’s t-test). **C.** Alizarin Red staining of wild-type sibling and *armc9^sa29834^* mutant fish at 5 wpf, visualized under stereomicroscopy at low (top row) and high (bottom row) magnification (mag). No vertebral bone fusion was observed in mutants, consistent with an idiopathic-like scoliosis phenotype. **D.** Scatter plot of number of vertebrae counted along the anterior-to-posterior axis of 5 and 8 wpf wild-type sibling and *armc9^sa29834^* fish. The average number of vertebrae in mutants was comparable to wild type (WT= 31.3, *armc9*= 30.57) at both post-embryonic stages. Sample sizes: wild type (n= 9 at 5 wpf and n= 7 at 8 wpf) and *armc9* (n= 6 at 5 wpf and n= 3 at 8 wpf). Data are presented as mean ± SEM. *p*=0.0023 (two-way ANOVA test). ns, not significant. **E.** Wild-type siblings and mutant larval development from 2 to 5 wpf, showing that the curved phenotype became apparent in *armc9^sa29834^* from 3 wpf onward, particularly in the caudal region. The alteration in the VC appears to hinder larval growth, as mutants were smaller than the wild-type larvae. **F.** Scatter plot of Cobb angle measurements in the caudal region of Alzarin Red-stained 5 wpf wild-type sibling and mutant fish. Vertebral curvature was significantly increased in mutants compared to wild-type individuals (WT= 0°, *armc9*=40°). Sample sizes: wild type (n= 11 at 5wpf) and n= 6 at 8wpf) and *armc9* (n= 8 at 5wpf and n= 3 at 8wpf). Data are presented as mean ± SEM. \*\*\*\**p*<0.0001 (two-way ANOVA test). wpf, weeks post-fertilization. Scale bar= 2 mm (A), 0.5 μm (C) and 0.75 μm (E).

To further investigate skeletal anomalies in *armc9* mutant fish, both adult wild-type and mutant individuals were stained with Alizarin Red at five weeks post-fertilization (wpf). In wild-type fish, the VC exhibited well-defined, continuous staining patterns indicative of normal bone formation and integrity (Fig. 1C). Bone structure analysis showed that both wild-type and mutant individuals displayed normal vertebral characteristics, including the typical biconic shape and the formation of hemal and ventral arches (Fig. 1C). These structural features were preserved even in regions affected by dorsal spine curvature (Fig. 1C). However, in *armc9* mutants, while the overall vertebrate structure was present, spinal bone fusion, commonly associated with skeletal abnormalities, was notably absent (Fig. 1C). Despite the severe spinal curvature, we did not observe incorrect vertebral segregation, even at the later developmental stages. Both wild-type and *armc9* mutant fish consistently exhibited between 30 and 32 vertebrae of similar size (Fig. 1D), aligning with the expected vertebral count in zebrafish (35). The absence of fused vertebrae in mutants supports the hypothesis that the scoliosis observed in these fish is idiopathic in nature, resembling human AIS, where the precise cause of spinal curvature remains unknown.

To determine the onset and progression of axial deformities, we monitored the development of wild-type and *armc9* mutant larvae from 2 to 5 wpf. Aberrant spinal curvatures first became apparent in the caudal region of mutant individuals at 3 wpf (Fig. 1D). This early onset of curvature was unexpected, suggesting that underlying skeletal abnormalities develop earlier than previously thought. As mutant larvae continued to grow, these curvatures become more pronounced along the dorsal-ventral axis, affecting overall body structure and alignment. In wild-type larvae, the spinal axis remained straight (0° curvature), whereas in *armc9* mutants, the average curvature reached 40° at 5 wpf and nearly 50° at 8 wpf, indicating progressive increase in severity (Fig. 1F). The reduced growth rate observed in mutants suggests that axial curvature may not only result from skeletal malformations but could also impair overall development, potentially disrupting essential physiological processes required for normal growth. These findings highlight a critical developmental window during which axial deformities emerge, underscoring their broader impact on larval growth and skeletal integrity.

### An inflammatory response is associated with the armc9 mutation

For years, the physiological processes underlying the development of early idiopathic scoliosis (EIS) have remained unclear. However, recent evidence suggests that inflammation may play a key role. In their study of *ptk7* mutants, Van Gennip and colleagues (2018) demonstrated that focal activation of pro-inflammatory signals within the VC induces spinal curvature. Additionally, macrophages were observed accumulating in the curved regions (23).

To assess tissue alterations adjacent to aberrant spinal curvatures in *armc9* mutants, we analyzed histological sections from both wild-type and mutant fish at 40 days post-fertilization (dpf). Our observations revealed tissue displacement caused by spinal curvature, with disorganized connective tissue, consisting of sparse extracellular matrix and scattered collagen fibers, invading spaces normally occupied by vertebrae (Fig. 2A). This displacement affected surrounding tissues, creating localized depressions. Notably, some mutant fish exhibited inflammatory foci, presumably composed of neutrophils and macrophages (Fig. 2A). In these areas, we also detected the formation of fibrotic connective tissue, predominantly surrounding the dorsal spine at the level corresponding to the loose connective tissue layer found between muscles and vertebrae in wild-type fish. Furthermore, we observed immune cell presence through the dorsal spine, with some cells infiltrating the notochord (Fig. 2A). The accumulation of immune cells in regions of spinal curvature suggests an active immune response, potentially indicating inflammation or tissue damage associated with scoliosis. The presence of these immune cells may contribute to the pathogenesis of the scoliosis phenotype in *armc9* mutants.

**Figure 2.**
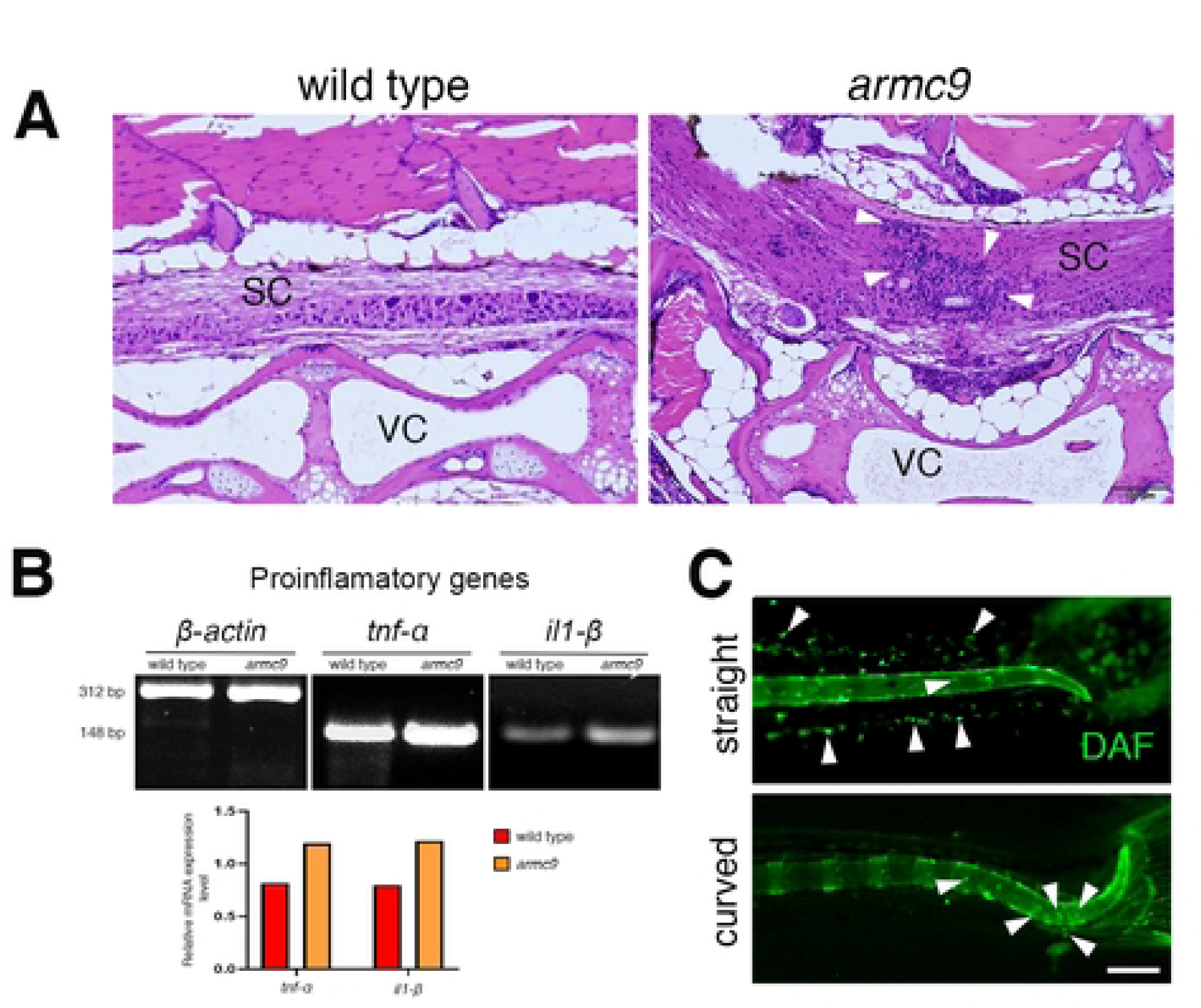
An inflammatory response is associated with spinal curvature. **A.** Histological sections of the caudal region in 40 dpf wild-type and *armc9^sa29834^* mutant individuals, stained with Hematoxylin-Eosin and visualized under optical microscopy. The wild-type siblings section shows a normal vertebral column (VC) with a straight spinal cord (SC). In contrast, *armc9* mutants display multifocal inflammation of the SC, leading to axial axis misalignment. A high magnification view, reveals an accumulation of presumably immune cells around the SC (white arrowheads). **B.** Top row: RT-PCR analysis of inflammatory markers, *tnfα* (148 bp) and *il1β* (149 bp), in 10 dpf wild-type and *armc9* larvae. *β-actin* is used as loading control. Bottom row: The increased amount of *tnfα* and *il1β* mRNA levels relative to *β-actin* indicate an excessive inflammatory response in *armc9* mutants. The relative expression levels of *tnfα* and *il1β* represent the intensity ratio of corresponding PCR product bands. **C.** Representative fluorescence images of 5 wpf straight and curved sibling larvae incubated with the NO-sensitive fluorescent sensor DAF-FM. In the straight larvae, NO-producing macrophages (white arrowheads) are observed along the caudal region. In the curved larvae, the accumulation of macrophages (white arrowheads) suggest that an elevated NO production is associated with the inflammatory state at the site of scoliosis onset, thus linking inflammation to the structural defect observed in *armc9^sa29834^* mutant fish. dpf, days post-fertilization; wpf, weeks post-fertilization; bp, base pairs; SC, spinal cord; VC, vertebral column, NO, nitric oxide. Scale bar= 100 μm (A) and 0.5 μm (C).

To further investigate the inflammatory response in mutant larvae, we analyzed the expression of key pro-inflammatory genes, *tnf-α* and *il1-β*, as markers of immune activation. Our results showed an upregulation of both genes in *armc9* mutants (Fig. 2B). Notably, this increased expression was more pronounced in the caudal region, coinciding with an accumulation of presumably immune cells (Fig. 2C). The presence of these cells and the elevated expression of inflammatory markers in curvature-prone regions suggest an active neuroinflammatory response associated with scoliosis development in *armc9* mutants.

To further explore the role of macrophages in scoliosis onset, we examined nitric oxide (NO) production, a key signaling molecule frequently associated with inflammation (36–39). Using the NO-sensitive fluorescent probe DAF-FM, we incubated 5 wpf straight and curved larvae to visualize NO-producing macrophages at scoliosis onset sites. Our results revealed a significant increase in DAF-FM fluorescence in the caudal region of curved larvae compared to straight ones (Fig. 2C), indicating elevated NO production. This suggests inducible Nitric Oxide Synthase (iNOS) activity and a heightened inflammatory state correlating with scoliosis onset (40, 41). These findings support a role for NO-producing macrophages in scoliosis development, directly linking inflammation to the structural abnormalities observed in *armc9* mutants.

### RF formation remains unaltered in armc9 mutant larvae

Recent studies on zebrafish mutants exhibiting axial curvature resembling scoliosis, such as *ptk7*, *scospondin*, and *dmh4* mutants, have identified defects in either CSF flow dynamics or in the structure of one of its major components, the RF (16, 19, 22). This fiber is formed by the aggregation of SCO-spondin protein, which is secreted by the SCO and, at early developmental stages, also by the flexural organ.

To investigate potential morphological alterations in the RF and SCO of *armc9* mutants, we performed immunodetection of SCO-spondin protein in larvae at 3 and 6 dpf individuals. At 3 dpf, the morphology of the SCO, flexural organ, and RF appeared comparable between wild-type and *armc9* mutant larvae (Fig. 3A). By 6 dpf, the SCO in wild-type larvae exhibited a funnel-shaped structure, with its posterior region giving rise to a single strand corresponding to the RF (Fig. 3B). Similarly, in *armc9* mutant larvae, no evident morphological defects were observed in the SCO or RF along the axial axis (Fig. 3B).

**Figure 3.**
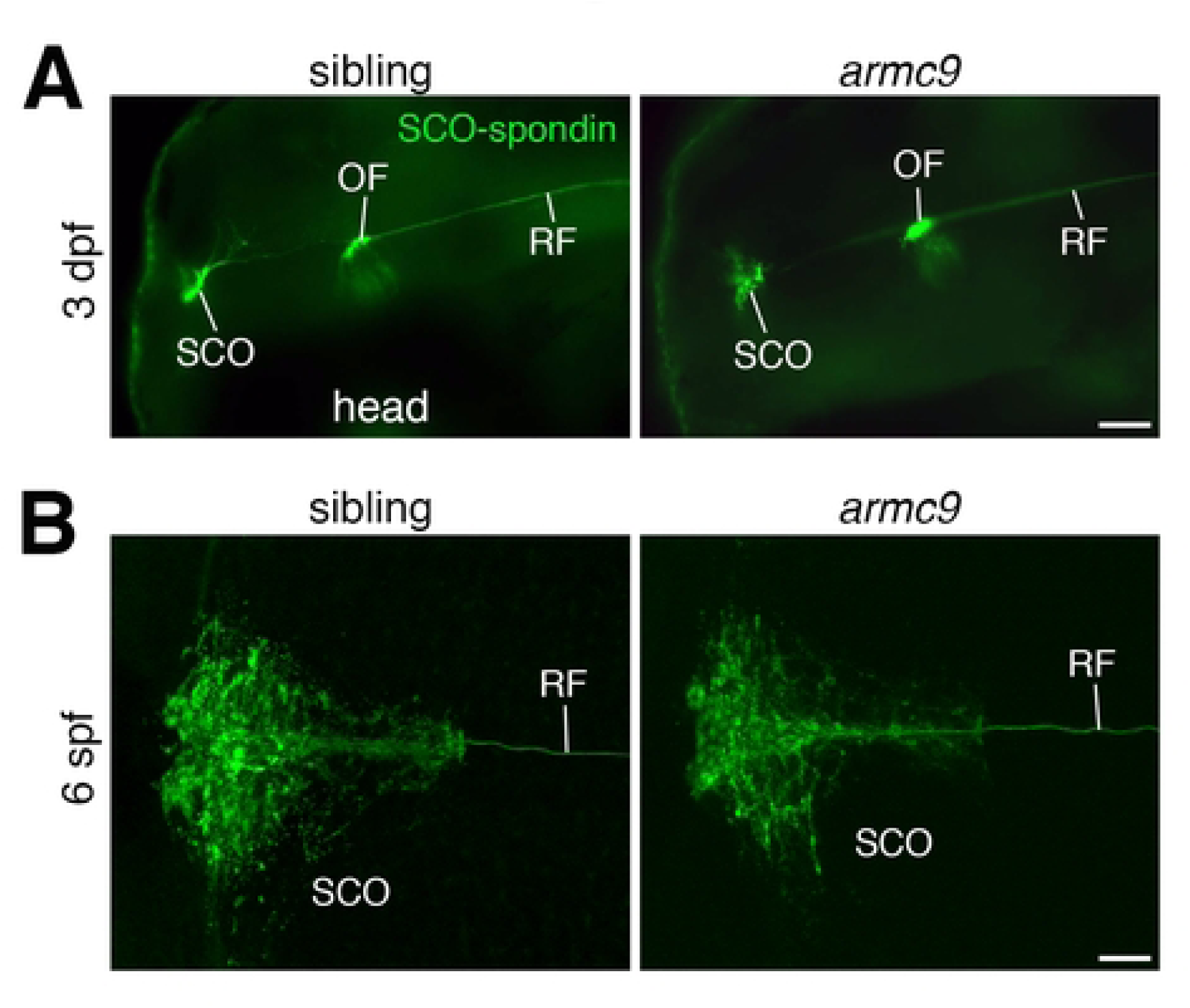
Morphology of the subcommissural organ in wild-type and *armc9^sa29834^* larvae. **A.** Immunodetection of the SCO-spondin protein reveals the morphology of the subcommissural organ (SCO), the flexural organ (FO), and the Reissner fiber (RF) in 3 dpf wild-type (n=6) and *armc9* (n=5) larvae head. **B.** Magnified confocal Z-projection view of the SCO in 6 dpf wild-type and *armc9* mutant larvae. No apparent differences in the SCO morphology were observed between wild-type (n= 5) and mutant (n= 3) individuals. dpf, days post-fertilization. Scale bar= 60 μm (A) and 20 μm (B).

Although SCO-spondin plays a well-stablished role in maintaining RF integrity and is critical for neurodevelopmental processes, our observations revealed no apparent morphological differences in SCO or RF structure among wild-type, heterozygous, and *armc9* mutant larvae. These findings suggest that the axial curvature and scoliosis observed in *armc9* mutants arise independently of major disruptions in SCO or RF morphology.

### The armc9 mutation in zebrafish leads to structural disruption

The *armc9* gene mutation was identified through whole-exome sequencing of sperm from mutagenized males (34). This gene spans 89.69 kilobases (kb), contains 18 coding exons, and encodes the Armadillo (ARM) repeat-containing protein 9 (Armc9) (Fig. 4A). Armc9 plays a crucial role in various cellular processes, particularly in protein-protein interactions mediated by its ARM-type fold domain and C-terminal region (31). The *sa29834* mutation specifically affects exon 16, where a cytosine (C) to thymine (T) transition introduces a premature stop codon in the *armc9* coding sequence (Fig. 4A). The wild-type Armc9 protein comprises of 817 residues, organized into multiple alpha-helical structures arranged in tandem with interconnecting coils. Its ARM-type fold domain (UniProt ID: IPR016024), spanning residues 383-578, is a key functional region implicated in protein interactions, highlighting the importance of this mid-segment in Armc9 function (Fig. 4B) (31, 42).

**Figure 4.**
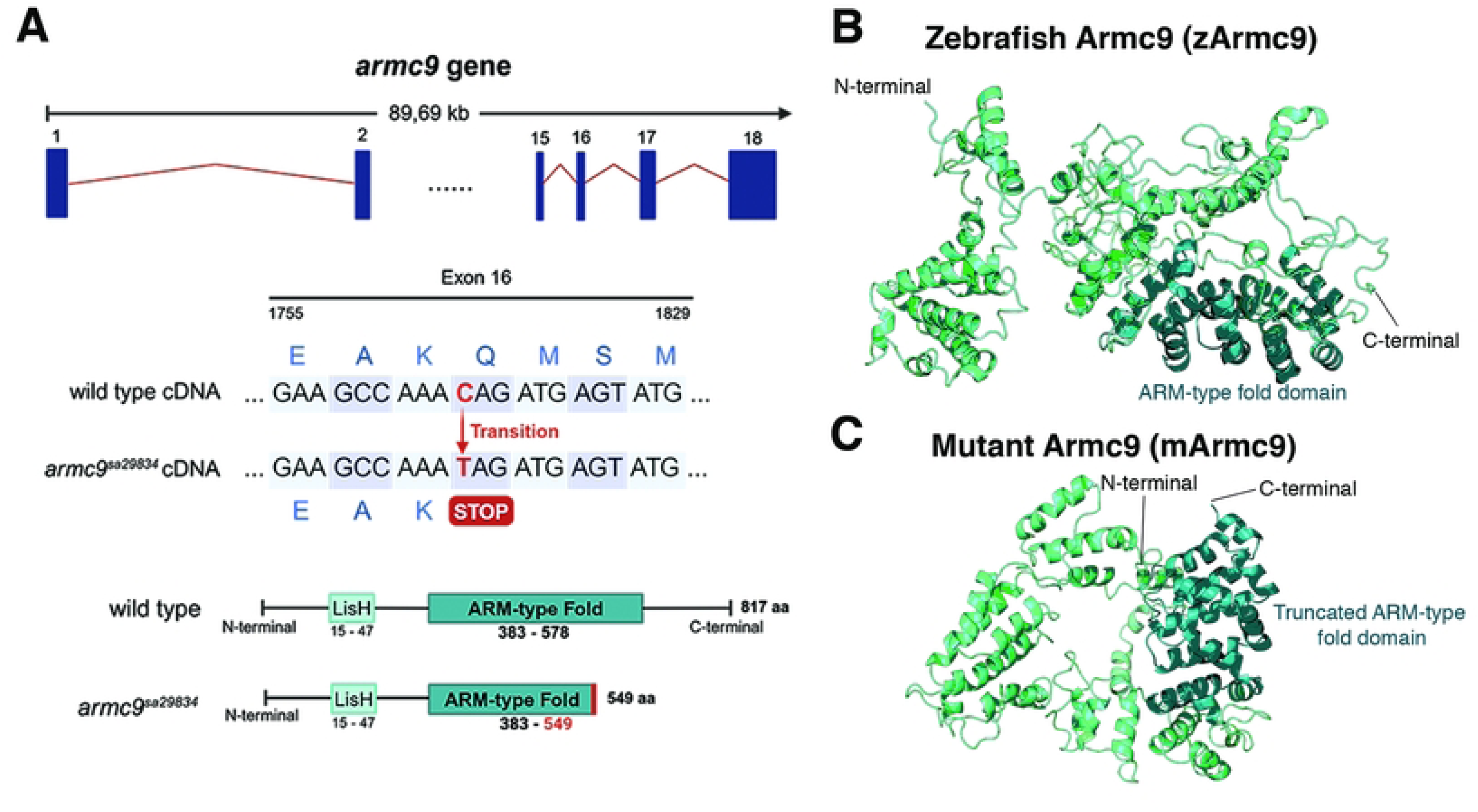
Impact of the *sa29834* point mutation on the zebrafish *armc9* coding sequence and protein structure. **A.** Top row: Schematic representation of the zebrafish *armc9* gene, which spans 89,69 kb and comprises 18 coding exons. Middle row: The *sa29834* mutation impacts on exon 16, provoking Cytosine (C) to Thymine (T) transition in the *armc9* and producing an early stop codon. Bottom row: Schematic representation of the *sa29834* mutation-effect at the protein level. The wild-type Armc9 protein consists of 817 amino acids, whereas the premature stop codon truncates the mutant protein at amino acid 549. This truncation leads to the loss of the ARM-type fold functional domain and remaining C-terminal portion. **B.** Tertiary 3D structural model of the zebrafish Armc9 (zArmc9) protein, composed of multiple alpha-helix structures arranged in tandem. **C.** Tertiary 3D structural model of the mutant Armc9 (mArmc9) protein, highlighting the loss of the C-terminal region, which alters its overall size and structural conformation.

As a result of the *sa29834* mutation, the truncated Armc9 protein comprises only 549 residues, leading to the loss of the C-terminal region and a significant portion of the ARM-type fold domain (Fig. 4A). This structural disruption likely compromises the protein’s functional capacity (Fig. 4B, C). Given the absence of an experimentally resolved three-dimensional (3D) structure for Armc9, we generated *in silico* models of both zebrafish wild-type and mutant proteins (Fig. 4B, C). As a result, the mutant Armc9 structure consists of only 549 residues due to the loss of the C-terminal portion, including a significant part of the ARM-type fold domain. This truncation alters the structural integrity and functional potential of the protein (Fig. 4B,C). Our results suggest a strictly post-embryonic role of zebrafish Armc9 controlling ciliogenesis and its activity mapped to the mid portion of the protein ((29), this work).

### Armc9 and Togaram1 form an interaction complex in zebrafish

Armc9 localizes to the proximal region of primary cilia in mammalian cells, although it has also been suggested to associate with the centriole (29, 30). This protein has been reported to interacts both directly and indirectly with multiple ciliary proteins, contributing to its interaction network (31). Among these interactions, ARMC9 directly binds CSPP1, CCDC66, and TOG1, the latter of which has been implicated in Joubert syndrome (31).

To investigate whether this interaction occurs in zebrafish and whether the axial curvature phenotype observed in *armc9* mutants could be attributed to an alteration in the Armc9-Tog1 complex, we performed an *in-silico* analysis of their interaction. The 3D structural models of zebrafish wild-type Armc9 (zArmc9), mutant Armc9 (mArmc9), and Togaram 1 (Tog1) were generated using AlphaFold2 (43), revealing distinct structural features associated with their respective sequences and functional domains. As described earlier, the *sa29834* mutation disrupts the structural integrity and functional potential of zArmc9 protein. Tog1, consisting of 587 residues (Fig. 5A), also contains an ARM-type fold domain between residues 348-548, which facilitates its interaction with zArmc9 (Fig. 5B, C).

**Figure 5.**
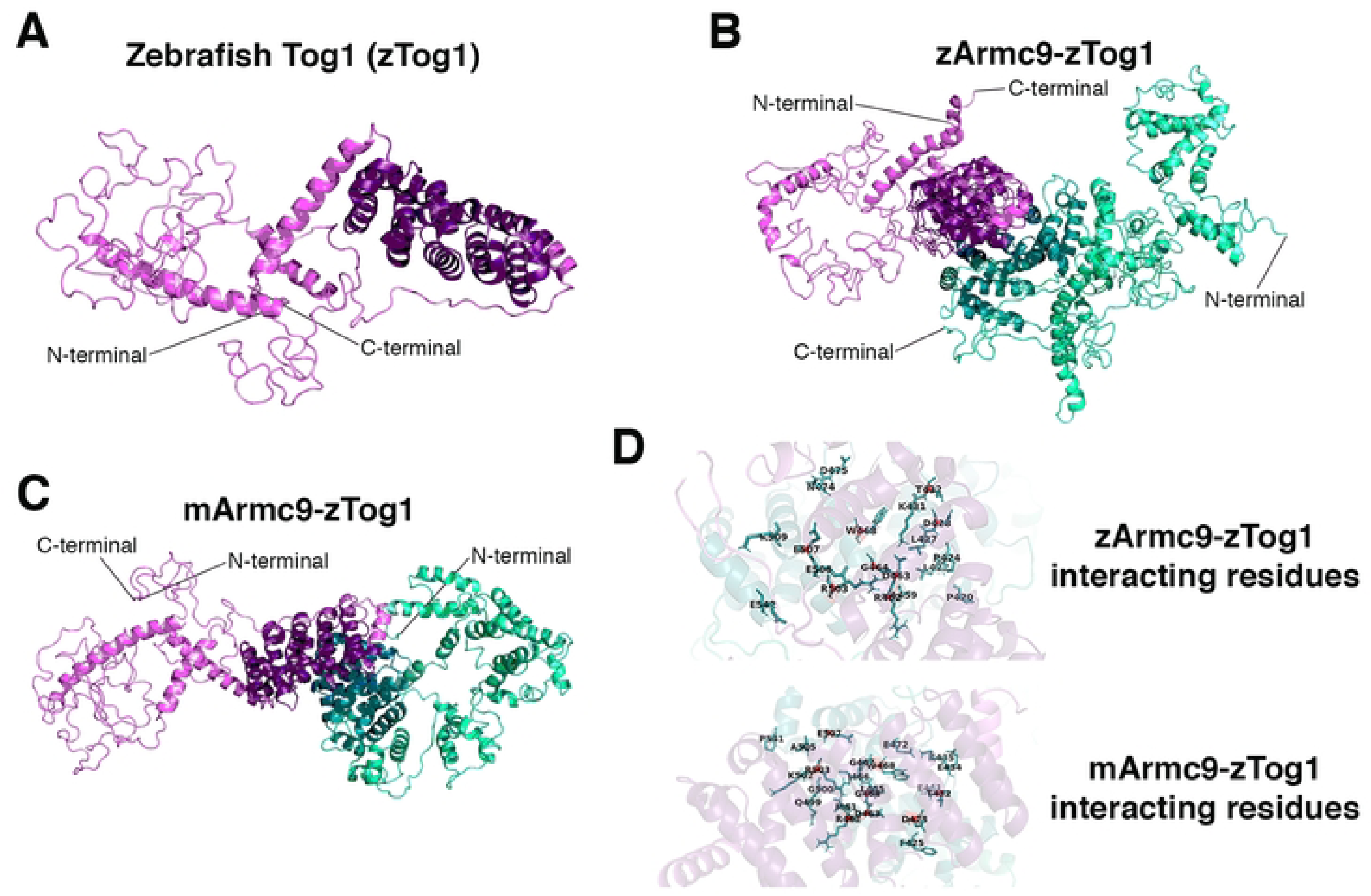
Interaction analysis of the zArmc9- and mArmc9-Tog1 complexes. **A.** Structural representation of zebrafish Togaram 1 (zTog1) (violet), including its respective ARM-type fold domain (deep teal and purple). **B.** Protein-protein interaction between the ARM-type fold domains of zArmc9 with zTog1, as predicted by HADDOCK simulation. **C.** Protein-protein interaction between the ARM-type fold domains of mArmc9 and zTog1. **D.** Top row: Interacting residues between zArmc9 and zTog1, highlighting key interface contacts. Bottom row: Interacting residues between mArmc9 and zTog1, illustrating differences due to the mutation. Shared residues involved in both interactions are marked in red.

To validate the generated 3D models, a comprehensive structural quality assessment was conducted using multiple computational tools. Global quality scores from QMEANDisCo (44) indicated moderate reliability of the models. ERRAT (45) confirmed acceptable atomic interaction accuracy, while ProSA Z-scores (46) placed the models within the range of high-quality protein structures derived from experimental data. Ramachandran plots generated via PROCHECK (47) showed minimal backbone geometry deviations. Additionally, PDBsum analysis (47) highlighted the secondary structure composition of the models, predominantly comprising helix-helix interactions (Supplementary Figure S1).

To further evaluate the structural stability and flexibility of the modeled proteins, molecular dynamics (MD) simulations were performed using GROMACS (48) for 100 ns. Structural deviations were assessed using Root Mean Square Deviation (RMSD) and Root Mean Square Fluctuation (RMSF) metrics (Supplementary Figure S2). Although RMSD values were higher than expected for compact globular proteins, they remained consistent with the flexible nature of the ARM-type fold domain and the presence of extended loop regions, which are known to mediate dynamic protein interactions (49). RMSF analysis revealed localized fluctuations, primarily in the N-terminal regions of each structure, corresponding predominantly to loop regions rather than the ARM-type fold domain. These findings support the functional and dynamic nature of the proteins and their suitability for subsequent docking analyses.

Following model validation, we performed molecular docking simulations using HADDOCK (50) and ClusPro (51) to investigate the interactions between zArmc9-Tog1 and mArmc9-Tog1, focusing on the ARM-type fold domains. The 25 residues with the highest contributions to binding free energy were ranked for each docking simulation (Table 1). In HADDOCK, the top-ranked zArmc9-Tog1 cluster exhibited a binding free energy of −124.89 kcal/mol, with 19 residues in the ARM-type fold domain contributing to the interaction. In contrast, mArmc9-Tog1 showed a weaker interaction, with a binding free energy of −64.33 kcal/mol and 22 contributing residues, reflecting the impacts of truncation. ClusPro results were consistent with those of HADDOCK, predicting zArmc9-Tog1 complexes with binding energies as low as −106.75 kcal/mol, while mArmc9-Tog1 interactions exhibited weaker affinities, reaching −85.66 kcal/mol (Fig. 5C, Table 1). HADDOCK identified eight conserved residues in chain A (zArmc9 and mArmc9) that interact with the ARM-type fold domain of Tog1 in both wild-type and mutant complexes. These residues, Arg462, Trp468, Asp463, Arg503, Glu507, Thr432, Asp428, and Gly464, are highlighted in red (Fig. 5D).

**Table 1.**
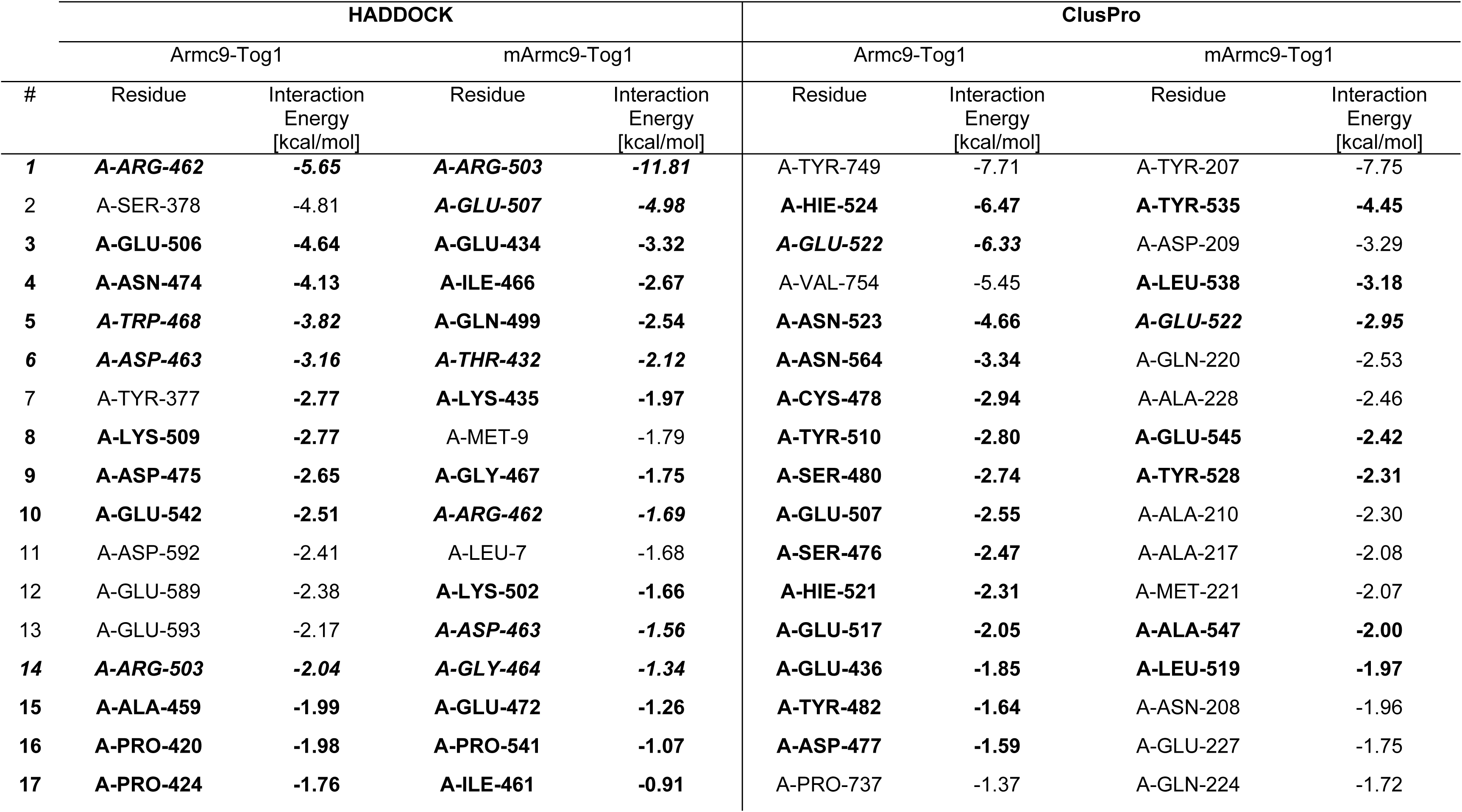
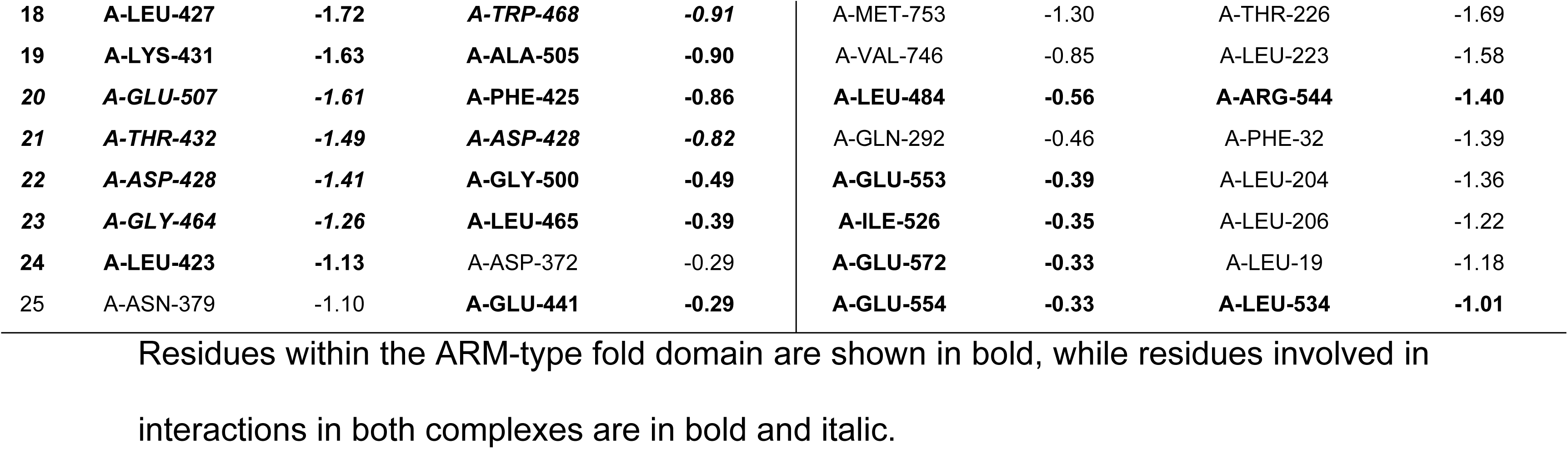
Top 25 ranked residues contributing to binding interaction energy in the zArmc9-Tog1 and mArmc9-Tog1 complexes, as identified through HADDOCK and ClusPro simulations.

Both docking simulations consistently indicated that the ARM-type fold domains play a crucial role in mediating interactions between Armc9 and Tog1. These findings suggest that the truncation of mArmc9 would reduce its binding affinity with Tog1, likely due to the loss of critical residues in the ARM-type fold domain.

### Evolutionary history of the ARMC protein family in animals

Armc9 belongs to the Armadillo repeat-containing (ARMC) protein family, which is characterized by the presence of an Armadillo (ARM)-type fold domain. A bioinformatic analysis of three model organisms, zebrafish, mouse, and human, identified 11 ARMC proteins in each species (Supplementary Table 1). Among these, two distinct functional domains were identified: the Armadillo repeat-containing domain (Arm-rpt) in Armc10 and the LisH domain in Armc9. Additionally, most ARMC proteins contained repeated elements classified as either Armadillo (Arm) or Arm-rpt, except for Armc1, Armc9, and Armc10, which lacked these features.

To investigate the phylogenetic relationships among ARMC proteins, we analyzed 12 species spanning major taxonomic groups of animals (Supplementary Table 2). All identified ARMC proteins shared a conserved Armadillo-like domain, classified as “ARM-type fold” (Table 2), making it a defining feature of the ARMC protein family. Using ARMC proteins sequences from humans, mice, and zebrafish, sequence similarity searches were performed with the BLASTp algorithm (52) against the coding sequences of the 12 species. This analysis identified a total of 148 homologous proteins, all of which contained the “ARM-type fold” domain, confirming their classification as ARMC family members (Supplementary Table 3).

**Table 2.** Functional domain analysis of ARMC family members in zebrafish.

| Protein/Domain | Domain |  | Homologous Superfamily |  | Repeat |  |  |
| --- | --- | --- | --- | --- | --- | --- | --- |
|  | InterPro |  | InterPro |  | InterPro | pfam | Prosite |
|  | Arm-rpt domain | LisH domain | ARM-like | ARM type fold | Armadillo | Arm | Arm-repeat |
| Armc1 | No | No | Yes | Yes | No | No | No |
| Armc2 | No | No | Yes | Yes | Yes | No | No |
| Armc3 | No | No | Yes | Yes | Yes | Yes | Yes |
| Armc4 | No | No | Yes | Yes | Yes | No | Yes |
| Armc5 | No | No | Yes | Yes | Yes | No | Yes |
| Armc6 | No | No | Yes | Yes | Yes | No | Yes |
| Armc7 | No | No | Yes | Yes | Yes | Yes | No |
| Armc8 | No | No | Yes | Yes | Yes | No | Yes |
| Armc9 | No | Yes | Yes | Yes | No | No | No |
| Armc10 | Yes | No | Yes | Yes | No | No | No |
| Armc1-like | No | No | Yes | Yes | Yes | Yes | No |

Several structural elements were identified among different ARMC proteins, with the ARM-like domain and the ARM-type fold being the only conserved features present in all ARMC proteins.

A multiple sequence alignment of these 148 proteins was then conducted, followed by the identification of phylogenetically informative sites and selection of the best-fit evolutionary model. A phylogenetic tree was constructed using the Maximum Likelihood method (Fig. 6), revealing 12 well-supported clades that included proteins from both vertebrate and invertebrate species. This broad taxonomic distribution suggests strong evolutionary conservation within the ARMC family. Notably, ARMC10, ARMC12, GPRASP, and ARMCX were grouped within the same clade, implying a close phylogenetic relationship and a likely shared ancestral origin. In the phylogenetic tree, ARMC9 clustered closely with ARMC8 and ARMC2, suggesting an evolutionary link between these proteins. The phylogenetic tree also showed high bootstrap values, indicating strong confidence in the inferred relationship (Fig. 6). All ARMC family members formed clusters with bootstrap values exceeding 80%, except for ARMC4, which had a lower but still acceptable support value of 57%. Despite this exception, the phylogenetic tree provides valuable insights into the evolutionary relationships among ARMC proteins.

**Figure 6.**
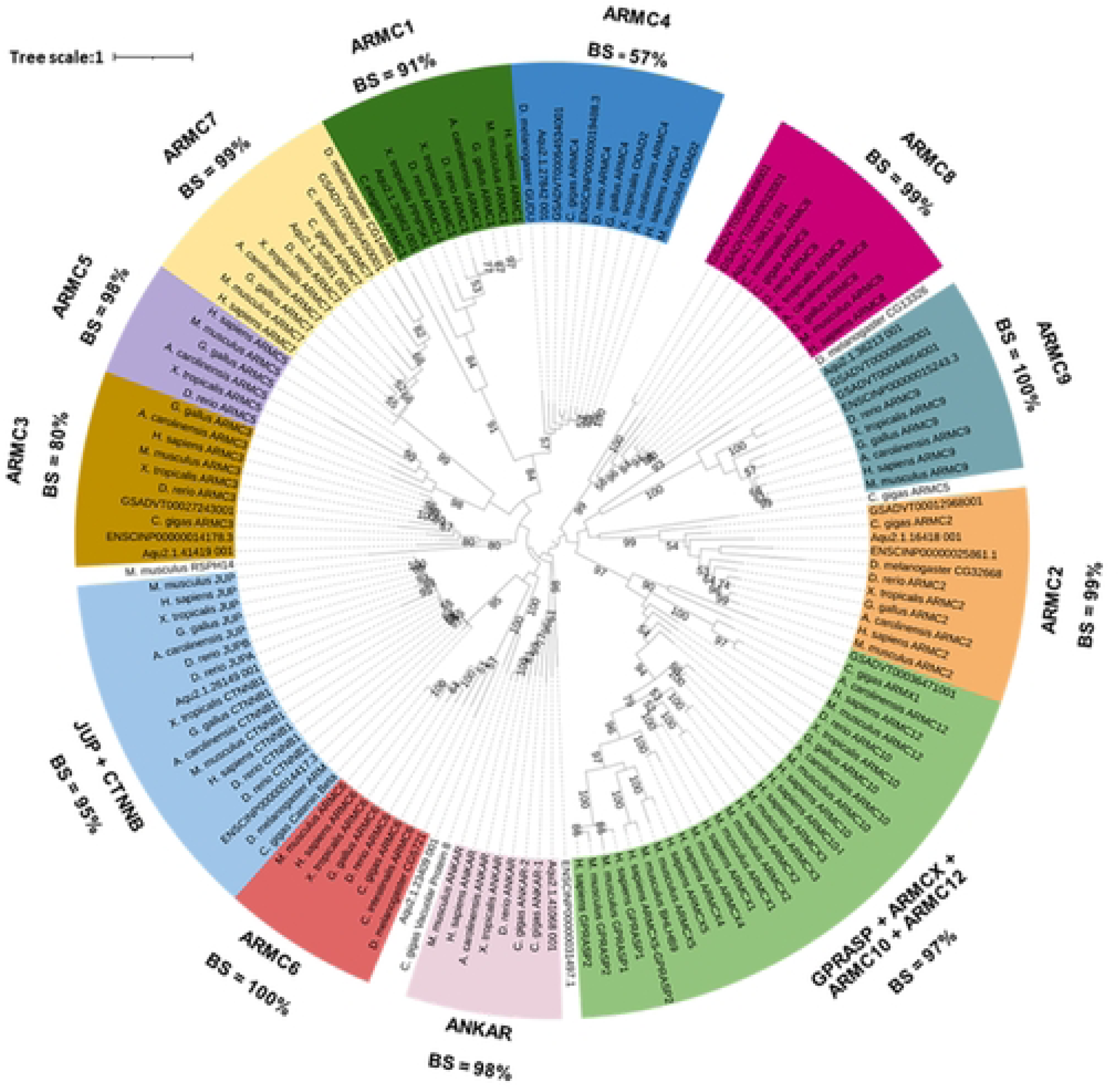
Phylogeny of ARMC family members in metazoans. Among the 148 proteins analyzed, 142 grouped into 12 distinct clades, while 6 remained unclassified. The presence of these 12 well-defined phylogenetic clades, encompassing both vertebrate and invertebrate species, suggests that ARMC family members have deep evolutionarily origins. Only bootstrap values >50 are shown. BS, bootstrap value.

Six proteins did not cluster within any of the 12 defined clades, suggesting that their sequences have diverged significantly, resulting in low sequence identity with homologs from other species. The ARMC9 clade contained proteins from both vertebrate and invertebrate species, including *Amphimedon queenslandica*, a marine sponge representative of the phylum Porifera. As sponges are considered the sister group to all animals (53), this finding suggests that the ARMC9 protein was already present in the common ancestor of metazoans. A similar pattern was observed in the clades of ARMC1, ARMC2, ARMC3, ARMC4, ARMC7, ARMC8, JUP + CTNNB, and ANKAR (Fig. 6). Conversely, the ARMC5 clade included only vertebrate species, suggesting that this protein originated in the common ancestor of vertebrates. Additionally, the clades containing ARMC6 and GPRASP + ARMCX + ARMC10 + ARMC12 comprised both vertebrates and invertebrates, indicating that these proteins likely emerged in the common ancestor of bilaterians.

Overall, these results indicate that ARMC family proteins appeared early in animal evolution, with their origins tracing back to the last common ancestor of metazoans.

## Discussion

AIS is a highly prevalent health condition with global significance (3), yet its etiology remains unknown. One of the main challenges in understanding the physiological processes underlying AIS is the lack of suitable animal models. Recently, the zebrafish model has provided significant advancements in elucidating the molecular pathology of AIS (5), offering crucial insights into its potential causes. In this study, we investigated the role of Armc9 in the establishment and maintenance VC in zebrafish. Through a combination of morphological and molecular phenotyping, bioinformatic strategies, and evolutionary analyses, we explored the etiology of scoliosis-like curvature in *armc9* mutants and examined the function of this genetic factor during the larva-to-juvenile transition. Additionally, we reconstructed the evolutionary history of ARMC family members across animal lineages. By integrating these findings, our study enhances the understanding of the genetic and molecular mechanisms involved in AIS development and highlights the regulatory role of *armc9* in this process. Unraveling these mechanisms is essential for identifying the underlying causes of AIS, a condition that significantly impacts the quality of life for millions of adolescents and adults worldwide.

### The zebrafish armc9 mutant as a model for AIS

Historically, various animal model systems have been employed to study scoliosis (5, 54). However, most of these studies have focused on inducing scoliosis-like curvature through mechanical or surgical means rather than investigating the molecular and cellular mechanisms underlying the development of this condition (55). With advancements in sequencing technologies, genetic models of scoliosis have become increasingly prominent, with zebrafish emerging as one of the most widely used organisms (54). Among these, the zebrafish *armc9* mutant has been proposed as a genetic model for studying AIS. This mutant was first described in the literature exhibiting hallmarks features such as retinal dystrophy, reduced cilia number and size, colobomas, and axial curvature (29). However, this initial characterization focused primarily on the adult phenotype, leaving many questions unanswered regarding the onset and biological processes involved in scoliosis development.

AIS is defined by two key characteristics that distinguish it from forms of scoliosis: 1) its specific timing of onset and 2) the absence of vertebral malformations (56). Unlike other zebrafish scoliosis models, the *armc9* mutant develops axial curvature during post-embryonic periods, specifically during the juvenile stage, rather than during embryonic development or adulthood. Between two and three weeks of development, curvature becomes apparent in the posterior axis, just before the caudal fin (Fig. 1). In contrast, other zebrafish scoliosis models, such as the *ptk7*, *zmynd10*, *kif3b*, and *sspodmh4/dmh4* mutants, exhibit axial curvature during embryogenesis (21, 22, 57). Thus, in terms of developmental timing, the onset of scoliosis in the *armc9* mutant more closely parallels AIS in humans.

Furthermore, axial curvature in *armc9* mutants arises independently of structural vertebral deformities, fusion, or changes in vertebral number, making it more analogous to human AIS. Notably, the absence of vertebral fusion in these mutants is particularly significant, as it supports an idiopathic-like scoliosis phenotype. This condition is characterized by lateral spinal curvature without apparent bone fusion defects, instead resulting from intrinsic vertebral misalignment and curvature (Fig. 1C). Understanding the timing and progression of these curvatures provides valuable insights into the role of *armc9* in vertebral and body axis development, with potential implications for developmental disorders. Given its developmental timeline and morphological characteristics, the *armc9* mutant represents a compelling model for studying the genetic and molecular basis of AIS.

### CSF flow remains unaltered in early developmental stages of the armc9 mutant

Aberrant CSF flow has been proposed as an early physiological mechanism contributing to scoliosis development (5). Notably, zebrafish *ptk7* mutant exhibit defects in CSF flow, drawing interest in both the dynamics and architecture of this system (16). One key structure within the CSF is the RF, a threadlike assembly formed by the aggregation of SCO-spondin secreted by the SCO (58). Zebrafish mutants lacking functional SCO-spondin either fail to develop or improperly assemble the RF, leading to defects in axial body straightening during embryogenesis (19, 20, 22). However, *armc9* mutants do not exhibit abnormalities in SCO morphology or significant defects in RF assembly (Fig. 3). Since proper RF formation depends on CSF flow (59), this suggests that CSF dynamics remain intact in *armc9* mutants.

Although studies have demonstrated a causal link between RF defects and scoliosis, not all zebrafish scoliosis models exhibit RF abnormalities. Indeed, other mutants, such as *kif7* or *adamts9*, develop axial curvature without RF defects during larval stages, similar to *armc9* fish (60, 61). This suggests that scoliosis associated with *armc9* mutations may be independent of both CSF flow and RF formation. Our assessment of SCO structure was conducted during early developmental stages, leaving open the possibility that structural alterations may emerge later. While the development of an AIS-like curvature in zebrafish appears compatible with normal SCO and RF formation, further experiments tracking RF dynamics throughout development, particularly in juvenile stages before curvature onset, are necessary to confirm its unaltered state relative to wild-type individuals. Additionally, given that RF function relies on CSF flow, direct assessment of flow dynamics are crucial, as they play a key role in axial patterning (16). Finally, since Armc9 is a ciliary protein, further investigation into cilia morphology within the central canal of the spinal cord is warranted. Establishing potential functional relationships between ciliary structure and CSF flow could provide deeper insights into the role of CSF components in axial curvature formation in the *armc9* mutants.

### Inflammation and macrophage recruitment in the development of axial curvature in the zebrafish armc9 mutant

Although the etiology of scoliosis remains unclear, dysregulation of inflammatory processes has been proposed as a contributing factor to axial curvature formation in both animal model systems and humans (reviewed in (5)). The link between immune system dysregulation and scoliosis development has also been proposed in zebrafish (62). For example, *stat3* loss-of-function mutants exhibit severe lateral and vertical spinal curvatures during in juvenile stages, with transcriptomic analysis revealing altered expression of genes involved in immunity and infection response (63). In another study, Van Gennip and colleagues (2018) demonstrated that focal activation of pro-inflammatory signals within the spinal cord is sufficient to induce a scoliosis-like phenotype in zebrafish. Notably, treatment with a non-steroidal anti-inflammatory drug mitigated curvature progression, further implicating inflammation in scoliosis pathogenesis (23). In *ptk7* mutants, an increased number of macrophages in the dorsal spinal region suggests an inflammatory response. However, in the *sspodmh4/+* mutant, despite elevated expression of the Tnf-α in the telencephalon of juvenile fish, macrophage recruitment was not observed in that region (22).

Our study provides compelling evidence that inflammation plays a role in the development of scoliosis in the zebrafish *armc9* mutant. Histological analysis revealed areas with tissue displacement around spinal curvatures. These regions contained localized inflammatory foci primarily composed of immune cells, presumably neutrophils and macrophages. This inflammatory response was further supported by the upregulation of *tnf-α* and *il1-β*, two key pro-inflammatory cytokines, along with increased NO production in mutant fish. These findings suggest that inflammation contributes to scoliosis pathogenesis, consistent with previous reports demonstrating that focal activation of pro-inflammatory signals within the spinal cord leads to curvature formation (23).

Future studies should investigate immune cell migration at different developmental stages and assess the expression of molecules related to the inflammatory response. Such analysis could clarify whether inflammation actively drives axial curvature formation in *armc9* mutants. Collectively, our results support a model in which inflammation, possibly triggered by tissue stress or mechanical disruption, contributes to scoliosis progression. Identifying upstream regulators of immune activation and testing whether modulating inflammation could mitigate spinal curvature in this model will be crucial next steps.

These findings shift the focus away from CSF composition or bone abnormalities as primary drivers of spinal deformities in *armc9* mutants. Instead, they highlight the potential role of alternative molecular and cellular mechanisms, particularly inflammation, in the onset of scoliosis. Further research is needed to explore these mechanisms, but our study provides a novel perspective on the factors contributing to spinal curvature in *armc9* mutants and potentially other contexts of spinal development disorders.

### Interaction between Armc9 and Tog1 depends on the ARM-type fold domain in zebrafish

Latour and colleagues (2020) identified Armc9 as a direct interactor of Tog1, CCDC66, and CSPP1 using an interaction screen with human Armc9 as bait. These interactions occur via the ARM-type fold domain. Notably, the interaction between Armc9 and Tog1 was mapped to amino acids 150-665, suggesting a functionally relevant protein-protein interaction region (31). However, this identified region spans nearly three-quarters of the total protein, making it relatively broad and imprecise. The ARM-type fold domain is known for its role in mediating protein-protein interactions. Its truncation due to the *sa29834* mutation likely disrupt Armc9’s ability to perform essential cellular functions. The resulting truncated protein underscores the importance of Armc9’s full-length structure for proper function. This mutation may contribute to developmental defects, including scoliosis observed in *armc9* mutants. However, the precise molecular mechanisms through which Armc9 regulates spinal curvature remain unresolved. Additionally, the loss of the C-terminal domain may further impair protein function, leading to a complete or partial loss-of-function. Such disruptions likely affect multiple pathways critical for normal development and cellular homeostasis.

Since experimentally determined 3D structures of zebrafish Armc9 and Tog1 are unavailable, we generated *in silico* models to conduct molecular docking analysis, using the AlphaFold2 server. Despite AlphaFold2’s advancements in structure prediction (43, 64), its models require thorough evaluation to ensure their suitability for downstream applications. Therefore, after generating 3D structures for wild-type and mutant Armc9, and Tog1, we refined the models using MD simulations with GROMACS to enhance their structural stability. The refined models were evaluated using QMEANDisCo, ERRAT, ProSA, and Ramachandran plots, confirming their quality for interaction studies. The ARM-type fold domain was clearly defined in all three structures, reinforcing its functional relevance. These refined AlphaFold2 models were subsequently used for docking analyses.

Rigid docking of wild-type and mutant Armc9 with Tog1 was performed using two widely recognized docking platforms: HADDOCK and ClusPro. Both platforms consistently identified the ARM-type fold domain as the primary interaction site between Armc9 and Tog1. Across all analyses, 25 amino acids with the lowest interaction energy were mapped to residues 383-578, significantly narrowing the interaction region to 195 amino acids, considerably more precise than the previously described in human ARMC9-TOG1 interaction site (31). In contrast, the mutant Armc9 exhibited weaker interactions with Tog1. The truncation of 29 residues within the ARM-type fold domain (residues 383-549) led to a substantial loss of key interaction sites, likely affecting the functional integrity of the Armc9-Tog1 complex.

To further validate the interaction between Armc9 and Tog1, we assessed the energetic parameters of the molecular docking models. Interaction energy, which measures binding stability, is a key indicator of interaction strength, with more negative values reflecting stronger and more stable interactions (65). Both HADDOCK and ClusPro predicted that the wild-type Armc9-Tog1 interaction was significantly more stable than that of the mutant complex. The truncation in *armc9* mutants led to a substantial reduction in interaction energy, suggesting a weakened binding potential due to the loss of key residues in the ARM-type fold domain. Loss-of-function mutations in Armc9 or Tog1 in human result in shorter cilia (31). Therefore, the scoliosis phenotype observed in *armc9* mutants may result from the disrupted cilia stability, ciliary signaling pathways, and function. Interestingly, HADDOCK analysis revealed that 8 out of the 25 lowest-energy interaction residues were conserved between wild-type and mutant Armc9-Tog complexes. This suggests that while some key residues remain functional, the overall interaction is compromised in the mutant.

Future experiments should focus on targeted mutagenesis of the amino acids identified as critical for the Armc9-Tog1 binding. Validating these residues through site-directed mutagenesis and biochemical assays will provide deeper insights into the molecular mechanisms governing this interaction. Additionally, investigating how the disruption of this complex affects ciliary signaling pathways may help elucidate the role of Armc9 in spine development and scoliosis pathogenesis.

### Insights into the evolution and functional implications of the ARMC protein family

The ARMC protein family consists of various members characterized by the presence of tandem repeats of approximately 42 amino acids each (66). While the functions of several ARMC proteins have been previously described (67), the roles of many others at the cellular level remain poorly understood. To better understand the evolutionary relationships among ARMC family members and infer their functional similarities, we constructed a molecular phylogeny based on amino acid sequences vertebrate and invertebrate representatives. All ARMC proteins analyzed from zebrafish, humans, and mice contain the ARM-type fold domain, alongside additional domains that distinguish them and contribute to their functional diversity (67). As a criterion for establishing phylogenetic relationships within the ARMC family, we focused on the ARM-type fold domain, excluding other domains that are present in certain members but not universally conserved. Previous phylogenetic studies performed by Gul and colleagues (2017), have analyzed Armadillo proteins using only the repeated ARM domain sequences. However, because not all proteins with an ARM-type fold domain contain these repeats, such an approach likely excluded some members of the ARMC family (68).

Our phylogenetic analysis yielded low confidence values in the internal branches, likely due to the relatively low sequence identity among the repeated Armadillo domains in the ARMC proteins (69, 70). Additionally, using full-length protein sequences increases sequence divergence, complicating the resolution of deeper phylogenetic relationships. However, higher statistical support values were observed in more external branches of the phylogenetic tree, leading to the identification 12 distinct clades. Within these clades, members of the ARMCN family clustered together, with “N” corresponding to ARMC1 trough ARMC9. Notably, ARMC10 and ARMC12 grouped with ARMCX and GPRASP proteins, a pattern consistent with previous findings that the ARMCX family originated via retrotransposition of the Armc10 gene on human chromosome 7 (71). The close proximity of ARMCX and ARMC10 in our phylogenetic tree aligns with this evolutionary history. Functionally, ARMCX1, ARMCX3, ARMC10, and ARMC12 are all regulated by the Wnt/β-catenin pathway and contribute to cell cycle regulation and mitochondrial dynamics (71–75). Their functional similarities further support their phylogenetic clustering in our analysis.

Among the ARMC protein families, only ARMC1 and ARMC4 showed strong statistical support for deriving from a common ancestor. Interestingly, although these proteins grouped together in our phylogenetic tree, no functional similarities have been described between them. ARMC1 has been reported to localize in both the cytosol and mitochondria, where it plays a role in mitochondrial distribution (76). In contrast, ARMC4 is involved in the assembly of the outer dynein arm in the primary cilia, and its loss of function is associated with primary ciliary dyskinesia (77). Structurally, neither ARMC1 nor ARMC4 has been experimentally resolved in three dimensions, and computational models have not revealed significant structural similarities between them (67). Further studies are needed to determine whether there is a functional or structural basis for their evolutionary relationship. Vertebrate ARMC9 proteins formed a well-supported clade alongside proteins from selected invertebrates, including rotifers, sponges, and urochordates. This phylogenetic distribution suggests that ARMC9 was already present in the common ancestor of animals that lived approximately 700 million years ago (78, 79). Data from the Ensembl database indicate that the four invertebrate proteins closely related to vertebrate ARMC9 proteins share both the ARM-type fold region and the LisH domain, highlighting a high degree of structural conservation over evolutionary time.

Overall, our phylogenetic analysis revealed that ARMC proteins cluster into 12 distinct clades, suggesting that nine of these clades, including ARMC9 clade, originated in the last common ancestor of animals. Future studies on ARMC proteins function will build on these evolutionary relationships. The use of model organisms, such as zebrafish, to generate new mutants will provide crucial insights into the biological roles of ARMC proteins and further refine our understanding of their phylogenetic classification.

## Conclusion

This study provides a comprehensive characterization of the zebrafish *armc9* mutant, integrating phenotypic and bioinformatic analyses to gain insights into AIS and the functional roles of Armc9 and the ARMC protein family. Our findings support the notion that the curved phenotype observed in *armc9* mutants is not attributable to vertebral malformations, thereby establishing this model as representative of AIS. Furthermore, we found no significant disruptions in the CSF flow, RF formation, or SCO development. These results suggest that AIS in *armc9* mutants may arise through alternative mechanisms beyond conventional vertebral abnormalities.

Our bioinformatic analysis identified the presence of LisH and ARM-type fold domains in the zebrafish Armc9 protein. Structural modeling predicted that the *armc9* mutant lacks a functional ARM-type fold domain, a loss that may contribute to the AIS-like phenotype. Additionally, *in silico* interaction analysis suggested that Armc9 interacts with Tog1 through the ARM-type fold domain, providing novel insights into the molecular mechanisms underlying AIS. Phylogenetic analysis of the ARMC protein family further revealed the presence of 12 distinct evolutionary groups in animals. This classification not only contextualize the biological role of Armc9 but also provides a framework for exploring the potential functions of other ARMC family members. Integrating these results, we propose that *armc9* mutants accurately model the AIS phenotype and that the loss of the C-terminal ARM-type fold domain in Armc9 is a key contributing factor. Furthermore, our study highlights the role of Armc9 in skeletogenesis and an inflammatory response, both of which are essential for establishing and maintaining VC integrity. These findings challenge the conventional view of AIS as solely arising from vertebral malformations and suggest alternative pathogenic mechanisms involving ciliary function and inflammatory processes.

In summary, this study advances our understanding of AIS by elucidating the functional significance of Armc9 and the ARMC family and by proposing alternative mechanisms underlying VC malformations. These findings pave the way for future research to further unravel the complex developmental and pathological processes governing AIS.

## Acknowledgments

We thank Dr. Andrea Stout at the Cell & Developmental Biology (CDB) Department, University of Pennsylvania, as well as Germán Osorio at the CMA Bio-Bio, University of Concepcion, for their technical assistance with confocal microscopy. We also acknowledge the fish facility staff at the University of Pennsylvania, along with Andrea Aguilar, Karina Vega-Drake, and Ruth Cisternas at the University of Concepcion for their valuable technical support.

## Materials and Methods

### Maintenance and management of wild-type and armc9 mutant fish

Adult zebrafish of the wild-type AB strain and *armc9* mutants were maintained at 28°C in aerated, deionized water supplemented with 60 mg/L of Instant Ocean, adjusted to pH 7.0. The fish were kept under a 14-hour light/10-hour dark photoperiod. Wild-type, heterozygous, and homozygous recessive *armc9* embryos were obtained through natural crosses between adult zebrafish and subsequently raised in 1X E3 medium (stock 10X: 5 mM NaCl, 0.17 mM KCl, 0.33 mM CaCl_2_, 0.33 mM MgSO_4_, pH 7.0) supplemented with 0.01% (w/v) methylene blue.

All animal experiments were conducted in accordance with the regulations set by the Bioethics and Biosafety Committee of the University of Concepción, as well as the Institutional Committees for the Care and Use of Animals at the University of Pennsylvania and the University of Chile.

### Genotyping of juvenile and adult zebrafish

To determine the genotypes of larval and adult zebrafish, individuals were anesthetized using a tricaine solution (MS-222, Sigma-Aldrich) at a concentration of 0.002 and 0.02 mg/ml, respectively. Genomic DNA (gDNA) was extracted using the HotSHOT method (80) from tail fin biopsies in both larvae and adults. Genotyping was performed using the KASPar system (KBiosciences) with allele-specific primers designed to distinguish wild-type and mutant sequences (Table 3).

**Table 3.**
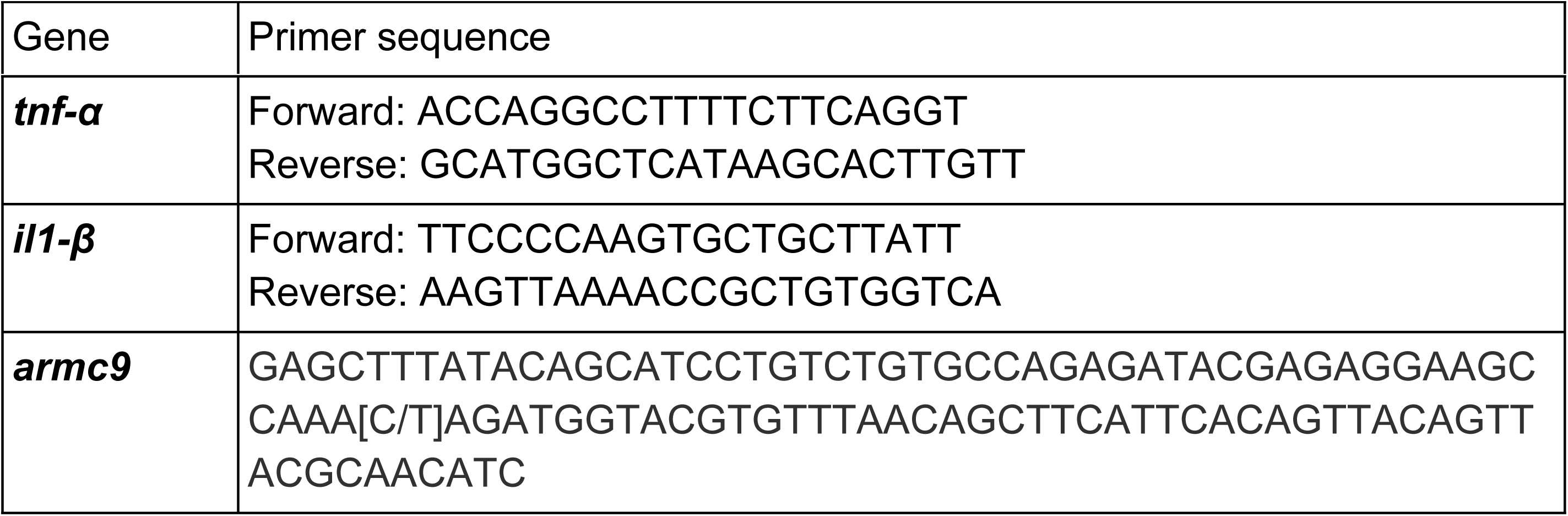
Primers sequences of pro-inflammatory markers and KASPar genotyping.

### Phenotyping of wild-type and armc9 mutant individuals

Phenotypic differences between wild-type and *armc9* mutant zebrafish were assessed at 2 and 8 wpf and in adulthood. The analyzed phenotypic traits included vertebrae count, curvature severity, and body length at each developmental stage. Images were taken using a Zeiss Stemi 508 equipped with an Axiocam 508 camera.

Statistical analyses were performed using a two-way ANOVA for vertebrae count and curvature severity, and a Student’s t-test for adult body length measurements, using GraphPad Prism v8.

### Cobb angle measurement

The Cobb angle, a standard parameter for assessing scoliosis severity in humans, was adapted for zebrafish larvae at 5 and 8 wpf. Curvature was defined using the central vertebra as reference point (designated as 0°), with two adjacent vertebrae on either side serving as angle vertices. Measurements were based on intervertebral angles, with the intersection point determining the curvature angle.

### RT-PCR analysis

Total RNA was extracted from 10-day post-fertilization (dpf) wild-type and *armc9* mutant larvae using Trizol-Chloroform, as described in (81). Complementary DNA (cDNA) was synthesized using the MMLV Reverse Transcriptase cDNA synthesis kit (GeneOn Cat# 105-100). Primers targeting pro-inflammatory markers genes, *tnf-α* and *il1-β,* were designed (Table 3). PCR amplification was performed using Maximo Taq DNA Polymerase 2x-preMix (GeneOn Cat# S113) in triplicate on 96-well plates with the following thermal cycling conditions: initial activation for 5 min at 95°C, followed by 45 cycles of 30 s at 95°C, 45 s at 60°C, and 35 s at 72°C. Gene expression levels were normalized to *β-actin* gene (*actb1*) gene. Relative mRNA expression levels (*tnfα*/*β-actin* and *il1β*/*β-actin* ratios) were determined by measuring the intensity of the corresponding PCR product bands using ImageJ software.

### Histology

To assess tissue integrity, 40 dpf zebrafish, both wild-type and *armc9* mutants, were euthanized using 160 mg/L tricaine (MS-222, Sigma-Aldrich) following a 24-hour fasting period. Specimens were then fixed in 10% neutral-buffered formalin at 4 °C for 24 hours, rinsed with 1X PBS, and decalcified in 0.35 M EDTA (pH 7.8) at room temperature for five days. After decalcification, samples were dehydrated and embedded in paraffin, following the protocol described in (82). Tissue sections were stained with hematoxylin and eosin for microscopic examination.

### Immunofluorescence and imaging

For SCO-spondin immunostaining, 6 dpf wild-type and mutant embryos were fixed in Carnoy’s solution (6:3:1 ethanol: chloroform: acetic acid) for 3 hours at room temperature, followed by rehydration through a graded ethanol series. Pigments were removed using 3% H_2_O_2_-0.5% KOH. Samples were then blocked with 2% BSA in 1X PBS supplemented with 0.5% Triton and 1% DMSO for 2 hours. Primary antibody incubation was performed using anti-SCO-spondin (Afru-11) at 1:100 dilution for 48 hours at 4°C, followed by incubation with an Alexa-488 conjugated secondary antibody (1:500) overnight at room temperature. Imaging was conducted using a Zeiss LSM 710 and LSM 880 confocal microscope with Plan-Apochromat objectives (10x/0.45, 20x/0.8, and water-immersion 40x/1.2).

### Alizarin Red staining of the zebrafish skeleton

Juvenile wild-type and *armc9* mutant zebrafish (5-8 wpf) were euthanized using tricaine (0.02 mg/ml) and fixed in 4% paraformaldehyde overnight at 4°C. Specimens were depigmented using a gradient of 1% KOH and 3% H_2_O_2_ in ethanol. Skeleton staining was performed with 0.5% Alizarin Red in 1% KOH at room temperature overnight. Vertebrae were manually cleaned of scales, and specimens were gradually transferred to increasing glycerol concentrations (25%, 50%, 75%, and 100%) in 1% KOH for storage. Imaging was conducted using a Zeiss Stemi 508 equipped with an Axiocam 508 camera.

### Nitric oxide detection

To detect nitric oxide (NO) along the anterior-posterior axis, straight and curved fish at 5 wpf were incubated in 5 μM 4-amino-5-methylamino-2′,7′-difluorofluorescein diacetate (DAF-FM, Sigma) for 3 hours at 28°C in darkness. Following incubation, larvae were rinsed in E3 medium and anesthetized with 0.002 mg/ml tricaine (MS-222, Sigma-Aldrich) for imaging. Fluorescence was visualized using a ZEISS Axio Zoom.V16 stereomicroscope.

### Molecular modeling of zArmc9, mArmc9, and Tog1 proteins

The 3D structures of zebrafish zArmc9, mArmc9, and Tog1 were predicted using AlphaFold2 via ColabFold (43). Amino acid sequences for these proteins were retrieved from UniProt (Armc9, ID: E7F187) and NCBI (Tog1, ID: XP_021322924.1). AlphaFold2 was optimized for high-performance modeling, applying six recycles for each protein. Multiple sequence alignments were generated using the mmseqs2_uniref_env mode, and post-modeling relaxation was performed using the AMBER protocol to resolve steric clashes and optimize model stability. Model confidence was assessed using pLDDT and pTM scores, ensuring accuracy at both local and global levels.

The resulting models underwent further validation using multiple structural assessment tools. QMEANDisCo (44) provided global quality scores to assess structural reliability, while ERRAT (45) evaluated non-bonded atomic interactions. Additional validation metrics included ProSA Z-scores (46) for overall model quality and Ramachandran plots generated via PROCHECK (47) to analyze backbone torsion angles. Residue-specific 3D/1D profiles were examined using Verify3D (83) to ensure structural compatibility with the primary sequences.

### Molecular dynamics simulations

To refine the structural predictions, molecular dynamics (MD) simulations were performed using GROMACS v2023.1 (48) on a computational cluster equipped with Intel Xeon Gold 6132 processors and NVIDIA Quadro P4000 GPUs. The highest-ranked AlphaFold2 models were embedded in orthorhombic SPC216 water boxes with a minimum padding of 1.0 nm. Counterions were added to neutralize the net charges and simulate physiological ionic strength. Energy minimization was carried out using the steepest descent algorithm, followed by equilibration under canonical (NVT) and isothermal-isobaric (NPT) conditions at 300 K and 1 atm, employing the V-rescale thermostat and Parrinello–Rahman barostat, respectively. The production phase was run for 100 ns, with frames recorded every 30 ps. RMSD analysis indicated structural stabilization within the first 20 ns, confirming the reliability of the models for further studies. The final stabilized frames were used for protein-protein interaction analyses.

### Protein-protein interaction analysis between zebrafish Armc9 and Tog1

To assess the molecular interaction between both zebrafish Armc9 and Tog1, docking simulations were performed using HADDOCK v2.4 (50) and ClusPro v2.0 (51). Structural models of zArmc9, mArmc9, and Tog1 were used as input for these simulations, with a specific focus on the ARM-type fold domain, identified as a potential interaction site. HADDOCK conducted semi-flexible docking with residue-specific restraints derived from the ARM-type fold domains, ranking clusters based on the lowest average interaction energy (65). In contrast, ClusPro employed rigid-body docking, prioritizing models according to cluster size, balanced score, and localization within the ARM-type fold domains (84). For both wild-type and mutant Armc9-Tog1 complexes, the selected docking models exhibited the most favorable binding free energy and the highest number of cluster members, ensuring alignment with predicted functional regions.

Binding free energies of the selected complexes were calculated using HawkDock (85), which applies MM/GBSA methods to quantify interaction strength. Docking results were visualized using PyMOL v3.0 (86), confirming the spatial compatibility and interaction localization within the ARM-type fold domains.

### Phylogenetic analysis of ARMC family members in animals

To investigate the phylogenetic relationships among proteins of the ARMC family in animals, amino acid sequences of ARMC family members from zebrafish (*Danio rerio*), human (*Homo sapiens*), and mouse (*Mus musculus*) were retrieved from the Uniprot database (87). For proteins with multiple isoforms, the canonical sequence, the longest variant, was downloaded (Supplementary Table 1). ARMC proteins from these three species were analyzed using the InterProScan server (Jones et al., 2014), revealing the presence of the ARM-type fold domain (InterProScan Domain: IPR016024) in all cases. When analyzing proteins from animal species listed in Supplementary Table 2, the presence of this domain was used as an exclusion criterion to determine which proteins would be included in subsequent phylogenetic analyses.

To identify homologous proteins within the ARMC family, BLASTP searches were performed (52). Only sequences containing the IPR016024 domain, as verified using InterProScan, were selected. The filtered sequences were then aligned using the MAFFT program with default parameters (88).

To retain phylogenetically informative regions, the multiple sequence alignment was processed with the trimAl program (89). The best-fitting evolutionary model for the phylogenetic analysis was determined using ProtTest3 (90). Phylogenetic reconstruction was conducted using the Maximum Likelihood method with RAxML program (91), and the resulting phylogenetic tree was visualized using iTOL (92).

## Supporting information

**Supplementary Table 1.** Genes of the ARMC family in zebrafish, human, and mouse, along with their corresponding UniProt accession codes.

| Gene name | UniProt code |  |  |
| --- | --- | --- | --- |
|  | Zebrafish | Human | Mouse |
| *armc1* | Q5PR58 | Q9NVT9 | Q9D7A8 |
| *armc2* | F1QBI9 | Q8NEN0 | Q3URY6 |
| *armc3* | E7F083 | Q5W041 | A2AU72 |
| *armc4* | E7FB07 | Q5T2S8 | B2RY50 |
| *armc5* | U3JA44 | Q96C12 | Q5EBP3 |
| *armc6* | A0A0G2K | Q6NXE6 | Q8BNU0 |
| *armc7* | F1R1C2 | Q9H6L4 | Q3UJZ3 |
| *armc8* | Q05AL1 | Q8IUR7 | Q9DBR3 |
| *armc9* | E7F187 | Q7Z3E5 | Q9D2I5 |
| *armc10* | F6NIY4 | Q8N2F6 | Q9D0L7 |
| *armc12* | - | Q5T9G4 | Q80X86 |
| *armc1-like* | F1R385 | - | - |

**Supplementary Table 2.**
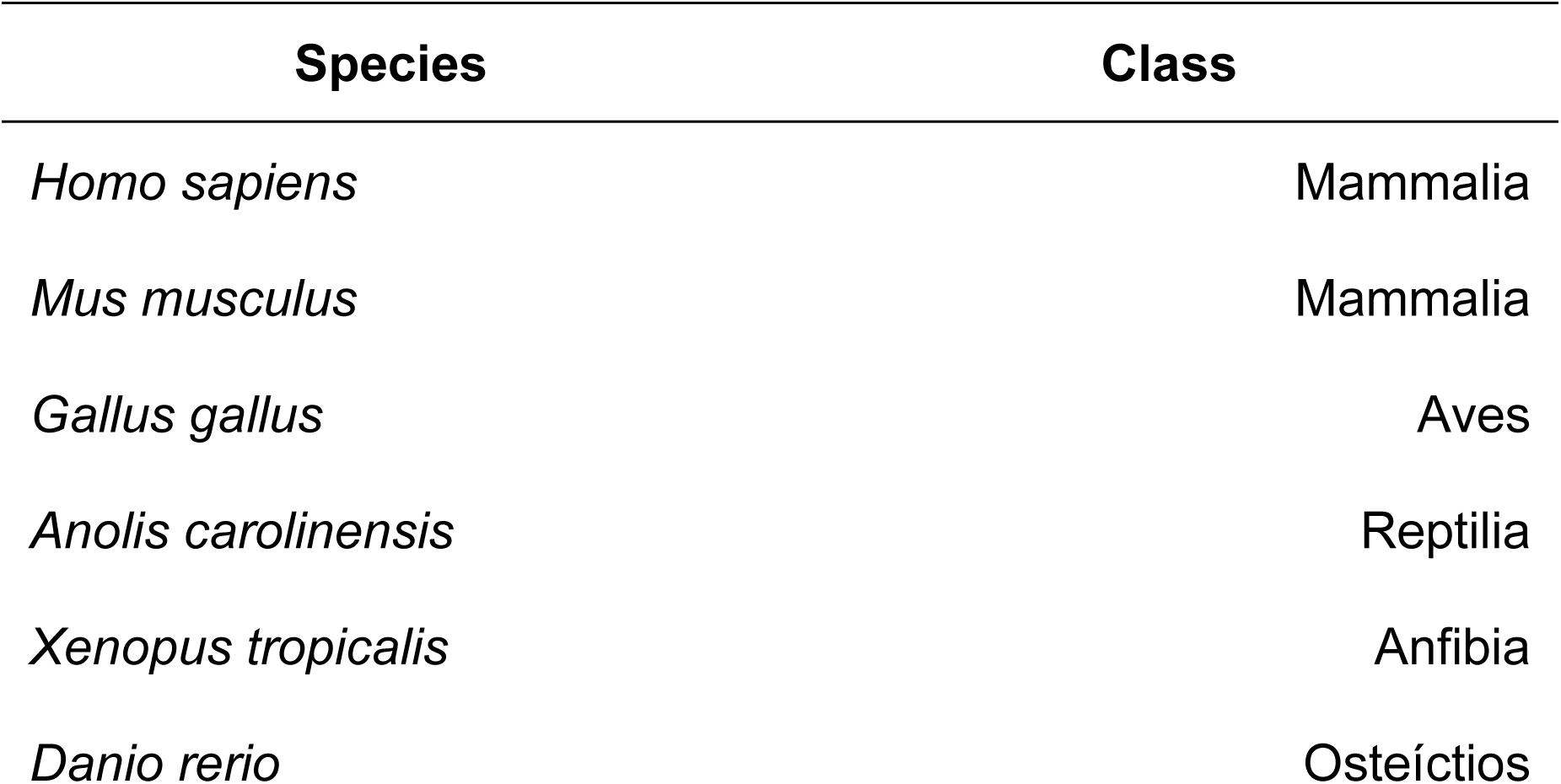
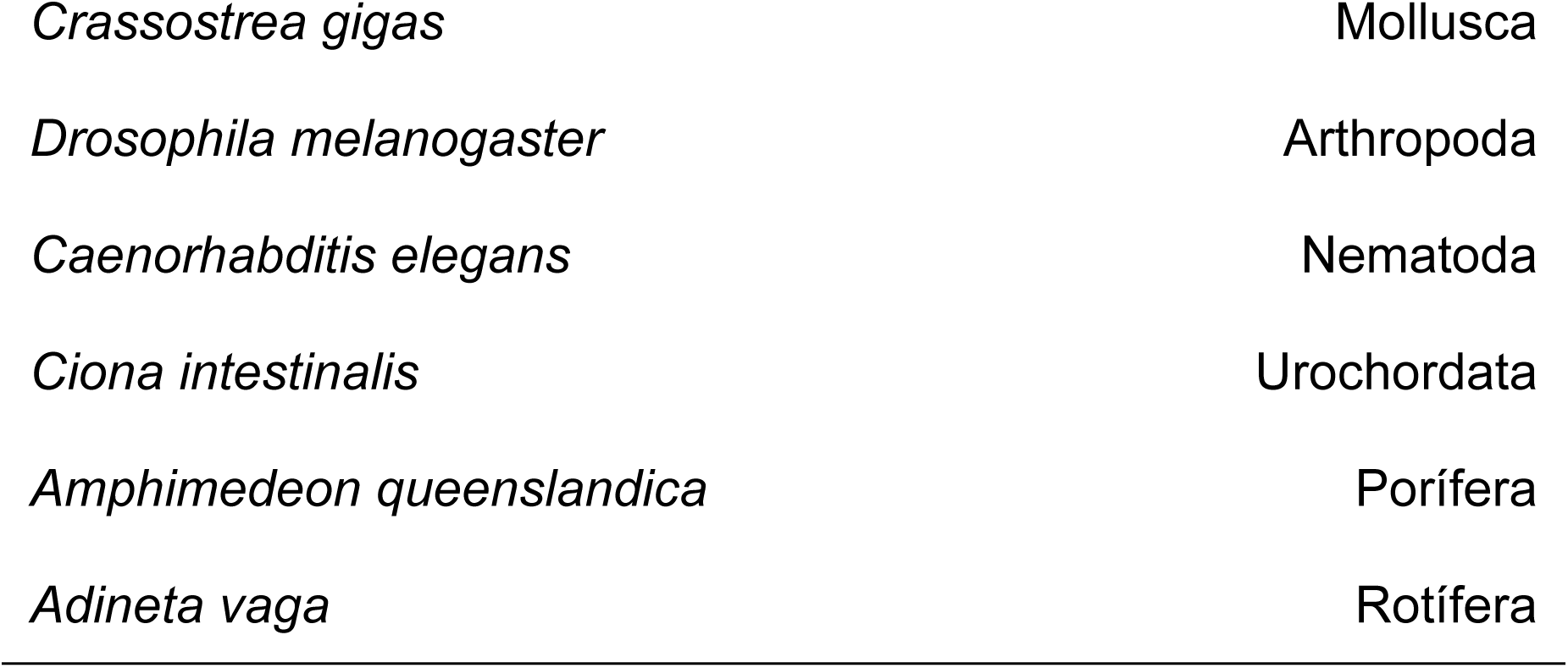
List of species included in the phylogenetic analyses. Protein sequences for these species were retrieved from the Ensembl database.

| Species | Class |
| --- | --- |
| *Homo sapiens* | Mammalia |
| *Mus musculus* | Mammalia |
| *Gallus gallus* | Aves |
| *Anolis carolinensis* | Reptilia |
| *Xenopus tropicalis* | Anfibia |
| *Danio rerio* | Osteictios |
| *Crassostrea gigas* | Mollusca |
| *Drosophila melanogaster* | Arthropoda |
| *Caenorhabditis elegans* | Nematoda |
| *Ciona intestinalis* | Urochordata |
| *Amphimedon queenslandica* | Porifera |
| *Adineta vaga* | Rotifera |

**Supplementary Table 3.** Number of proteins containing the ARM-type fold domain across species selected for phylogenetic analysis.

| Species | Number of proteins "ARM-type fold" |
| --- | --- |
| *Homo sapiens* (Mammalia) | 23 |
| *Mus musculus* (Mammalia) | 23 |
| *Gallus gallus* (Aves) | 12 |
| *Anolis carolinensis* (Reptilia) | 13 |
| *Xenopus tropicalis* (Amphibia) | 14 |
| *Danio rerio* (Osteictios) | 16 |
| *Crassostrea gigas* (Mollusca) | 12 |
| *Drosophila melanogaster* (Arthropoda) | 6 |
| *Caenorhabditis elegans* (Nematoda) | 1 |
| *Ciona intestinalis* (Urochordata) | 9 |
| *Amphimedon queenslandica* (Porifera) | 10 |
| *Adineta vaga* (Rotifera) | 9 |

